# Technologically-Assisted Communication Attenuates Inter-Brain Synchrony

**DOI:** 10.1101/2022.06.06.494185

**Authors:** Linoy Schwartz, Jonathan Levy, Yaara Endevelt-Shapira, Amir Djalovski, Olga Hayut, Guillaume Dumas, Ruth Feldman

## Abstract

The transition to technologically-assisted communication has permeated all facets of human social life; yet, its impact on the social brain is still unknown and the effects may be most notable during key developmental transitions. Applying a two-brain perspective, the current pre-registered study measured mother-child brain-to-brain synchrony using hyperscanning EEG at the transition to adolescence during live face-to-face interaction versus technologically-assisted remote communication. The live interaction elicited 9 significant cross-brain links between densely inter-connected frontal and temporal areas in the beta range [14-30 Hz]. Mother’s right frontal region connected with child’s right and left frontal, temporal, and central regions, suggesting its regulatory role in organizing the two-brain dynamics. In contrast, the remote interaction elicited only 1 significant cross-brain-cross-hemisphere link, attenuating the robust right-to-right-brain connectivity during live social moments that communicates socio-affective signals. Furthermore, while the level of social behavior was comparable between the two interactions, brain-behavior links emerged only during the live exchange, suggesting that remote interactions yield a somewhat thinner biobehavioral experience. Mother-child right temporal-temporal synchrony linked with moments of shared gaze and the degree of child engagement and empathic behavior was associated with right frontal-frontal synchrony. Our findings indicate that human co-presence is underpinned by specific neurobiological processes, suggest potential reasons for "zoom fatigue", and open a much-needed discussion on the cost of social technology for brain maturation, particularly among youth.

**Highlights:**

- Technologically-assisted communication is prevalent; yet, its impact on the social brain is unknown
- We measured mother-child brain-to-brain synchrony during live and technologically-assisted remote interaction
- The live interaction elicited 9 cross-brain links between densely inter-connected frontal and temporal areas in the beta range
- The remote interaction yielded only 1 significant cross-brain cross-hemisphere link
- Brain-behavior linked emerged only during the live interaction
- Further research should examine the cost of social technology to brain maturation, particularly among youth.

## 1. Introduction

From the moment infants are born, face-to-face communication is ubiquitous and supports the coordination of non-verbal social signals and the synchrony of physiological processes between parent and child (Feldman, 2017; Hari et al., 2015). Such bio-behavioral synchrony is a critical component of humans’ socio-emotional development and accompanies face-to-face interactions throughout life (Feldman, 2020; Mogan et al., 2017). Indeed, collaborative abilities, proficiency in reading others’ intent, and the capacity to share others’ mental states have been theorized as key determinants of humans’ supremacy over the animal kingdom (De Waal and Preston, 2017; Dunbar and Shultz, 2007). As socially-oriented creatures, humans’ daily social exchanges support the maturation of complex cognitive skills, empathic abilities, and brain structure and functions (Hari et al., 2015; Hasson et al., 2012).

One mechanism hypothesized to underpin the universal effects of face-focused communication is inter-brain synchrony (Babiloni and Astolfi, 2014; Czeszumski et al., 2020; Liu et al., 2018). Inter-brain synchrony is defined as the temporal coherence of neural dynamics between multiple brains and has become a growing focus of research in social neuroscience (Czeszumski et al., 2020; Liu et al., 2018; Reindl et al., 2018). Several features of face-to-face interactions have been highlighted as particularly important for enhancing inter-brain synchrony in ecological contexts, including the increased opportunities for shared gaze, social engagement, empathic resonance, and interpersonal reciprocity which are embedded in moments of social exchanges in daily life (Dikker et al., 2021; Djalovski et al., 2021).

Still, while our species’ enlarged brain has arguably expanded across primate evolution through social interactions in the natural ecology (Dunbar and Shultz, 2007), modern technology has offered a transfer of face-focused interactions to other modes of communication that do not require the partners’ physical co-presence, an evolution that stretches our cultural and biological heritage to uncharted territories. We now communicate remotely through a variety of platforms and social media channels, such as Zoom or Skype (Anderson and Jiang, 2018). Social communication via technology has become a daily practice not only with business associates but also within close relationships, generating a paradigm shift in the development of our species whose impact on the social brain is still unknown. With the COVID-19 pandemic, technologically-assisted communication became the main mode of social contact; children attended school via internet platforms, families met on Zoom, and classes, businesses, cultural activities, and, in fact, much of social life have turned into a technologically-assisted mode that enables people to keep in touch through screens. This shift may be especially hazardous for adolescents, whose need for social connections is paramount (Frost and Rickwood, 2017; Hoare et al., 2016), and are spending more time on screens than in live social interactions, with about a half of US teens reported being almost constantly online (Anderson and Jiang, 2018). The transition to adolescence is a period of rapid brain reorganization (Blakemore, 2008), rendering it a time of heightened vulnerability for psychopathology and increased risk for social maladjustment (Fuhrmann et al., 2015). As such, understanding how the shift to screen-mediated sociality impacts the social brain development of our youth is a key question for social neuroscience that bears critical implications for tomorrow’s world.

In the current study, we put this question to test by measuring brain-to-brain synchrony between mothers and their young adolescents during live face-to-face interaction versus technologically-assisted video chat. Researchers in social neuroscience have called to complement controlled experiments of the single brain with ecologically-valid studies of two-brain coordination during naturalistic social exchanges (Hasson et al., 2012; Redcay and Schilbach, 2019). It has been further argued that brain-to-brain synchrony is a key mechanism by which the mature brain externally-regulates the immature brain and tunes it to adaptive social life, particularly during sensitive periods in development (Feldman, 2015; Leong et al., 2017). We utilized hyperscanning EEG to pinpoint processes that sustain neural coordination when partners are co-present versus when they communicate remotely.

Inter-brain synchrony is impacted by the partners’ affiliation and tighter cross-brain coupling has been shown between affiliated partners compared to strangers, including romantic partners (Kinreich et al., 2017), close friends (Djalovski et al., 2021), therapists and clients (Zhang et al., 2018), and teacher-students (Bevilacqua et al., 2017). Across mammalian species, the mother-child bond is the context where processes of biobehavioral synchrony are first acquired and practiced (Feldman, 2020, 2012), and greater neural synchrony has indeed been found between children and their mothers as compared to a female stranger (Endevelt-shapira et al., 2021; Piazza et al., 2020; Reindl et al., 2018). We thus examined the effects of remote interaction on neural coordination within the mother-child relationship, the optimal context for such assessment where partners are familiar with each other’s non-verbal signals and can utilize even the partial cues available through screens. Adolescents are the first generation for whom technologically-assisted communication is natural and practiced daily, and this eliminates the potential confounding effects of unease or unfamiliarity on neural coordination.

Hyperscanning EEG studies have pinpointed several brain regions of inter-brain connectivity during naturalistic social interactions. These include; (a) homolog connectivity of same-area-same-hemisphere, such as temporal-to-temporal (Djalovski et al., 2021; Kinreich et al., 2017), central-to-central (Djalovski et al., 2021), and frontal-to-frontal connectivity (Azhari et al., 2019; Cui et al., 2012; Kruppa et al., 2021; Pan et al., 2017; Reindl et al., 2018; Wang et al., 2020); (b) cross-hemisphere same-region linkage, such as left temporal to right temporal connectivity; and (c) non-homolog multi-region linkage of same or different hemisphere, such as frontal-to-temporal (Pérez et al., 2017), frontal-to-parietal (Piva et al., 2017), central-to-temporal (Endevelt-shapira et al., 2021; Pérez et al., 2017), central-to-parieto-occipital and centro-parietal and parieto-occipital connectivity (Dumas et al., 2010). Notably, most studies reported right-hemisphere connectivity of homolog or non-homolog regions (Cui et al., 2012; Dumas et al., 2010; Endevelt-shapira et al., 2021; Jahng et al., 2017; Noah et al., 2020; Pan et al., 2017; Sciaraffa et al., 2021), suggesting that the right hemisphere, which matures early (Geschwind and Galaburda, 1985) and is implicated in non-verbal affective processing (Borod et al., 1998), may be particularly sensitive to two-brain communication. Such multiple areas of neural linkage underscore the richness of cross-brain possibilities afforded by naturalistic co-present interactions that may reflect distinct mechanisms triggered by different social goals.

Adolescence has rarely been studied from a two-brain perspective and our hypotheses were thus based on parent-child or affiliated adult pairs. Cross-brain studies of younger children and parents showed linkage mainly in frontal areas. In an fNIRS study of mother-child dyads (5-9 years) during cooperation versus competition, Reindl et al. (2018) found inter-brain synchrony in frontal areas, including dorsolateral prefrontal and frontopolar cortex during cooperation only (Reindl et al., 2018). Miller et al. (2019) replicated the frontal right dorsolateral and PFC synchrony during cooperation between mothers and their 8-13-year-old children (Miller et al., 2019). Wang et al. (2020) found that children (5-11) with ASD showed higher parent-child inter-brain synchrony in frontal regions during cooperation compared to non-interactive tasks and neural synchrony was modulated by autism symptoms, highlighting the contribution of the child’s ability for social engagement to cross-brain linkage (Wang et al., 2020). Finally, in cooperation versus competition tasks across wide age range (8-18), frontal-frontal neural synchrony emerged in both cooperation and competition (Kruppa et al., 2021), suggesting a shift in parent-child neural dynamics as children grow (Jager et al., 2015). Studies of other affiliative bonds, such as romantic partners or close friends, implicated temporal regions of cross-brain connectivity (Djalovski et al., 2021; Kinreich et al., 2017) and showed their links with episodes of shared gaze and the partners’ reciprocity and social engagement. Overall, these studies pinpointed frontal and temporal areas as potential targets for cross-brain linkage between attachment partners and demonstrated brain-behavior coupling with involved social behavior.

Framed within the emerging field of naturalistic cross-brain neuroscience, we formulated three hypotheses. First, we expected the live face-to-face interaction to trigger significantly more inter-brain connections across widely-distributed areas compared to the technologically-assisted communication. Second, focusing on frontal, temporal, and central regions that have been detected in prior hyperscanning EEG studies, we expected the live interaction to elicit inter-brain linkage of three types: (a) homologous (same area, same hemisphere), (b) cross-brain (same area, different hemisphere), and (c) multi-dimensional (cross-region of both same and cross hemisphere). Consistent with prior studies of affiliated partners, we hypothesized strong linkage of frontal and temporal regions. Finally, guided by the *bio-behavioral synchrony* frame (Feldman, 2021, 2017, 2012), we complemented assessment of neural synchrony with both micro-level coding and global rating of social behavior and focused on episodes of shared gaze and children’s empathic social engagement, in line with research pointing to their contribution to inter-brain synchrony (Dikker et al., 2021; Kinreich et al., 2017). Based on prior work (Djalovski et al., 2021; Endevelt-shapira et al., 2021; Kinreich et al., 2017; Levy et al., 2017), we expected brain-behavior coupling during the live social interaction (Mu et al., 2016), with more neural connectivity linked with increased gaze synchrony and greater social engagement but the degree of brain-behavior coupling in the remote interaction remained an open question. Finally, beta rhythms have been implicated in parent-child attachment processes in both mothers’ (Hernández-González et al., 2016; Kringelbach et al., 2008) and young adolescents’ brains (Pratt et al., 2018). In naturalistic cross-brain studies, beta synchrony has been shown to sustain communication between romantic couples and close friends (Djalovski et al., 2021), to underpin empathy and compassion (Ciaramidaro et al., 2018), and to link with behavioral social engagement and shared gaze (Dikker et al., 2021), and we thus focused our search on cross-brain beta-band synchrony.

## 2. Materials and Methods

The study was preregistered: https://osf.io/swun7/

### 2.1 Participants

We recruited 140 participants, comprising 70 mother-child pairs, through ads posted in schools and social media. Children were 12.26 years old (SD = 1.21, 44% males, 66% firstborn), healthy, and attended state-controlled typical (not special education) schools. Mothers were 43.74 years old (SD = 4.41), had an average 16.96 years of education (SD = 2.5), and were the biological mother and primary caregiver. All families were of middle-class background and 81% lived in the same household as the child’s father. The study was completed before the COVID-19 pandemic. The experiment was approved by the Reichman University institutional ethics committee and all mothers signed a written informed consent for themselves and their children. All procedures were explained to the participants prior to the experiment and they were free to leave the experiment at any time with full compensation. Participants were reimbursed for study participation ($30 per hour).

### 2.2 Procedure

The study took place in two adjacent experimental rooms and included three sessions recorded with dual-EEG. The first session was a baseline recording of mother and child’s brain (Rest) when partners are in the same room facing the wall and instructed not to interact. In the second session, the live interaction, mother and child were sitting facing each other and were instructed to socially interact on a planned positive topic (see below). In the third session, the video chat, mother and child communicated through a computer screen from two separate rooms and socially interacted on a planned positive topic with doors locked (Fig. 1A). All sessions were videotaped for offline behavioral coding.

**Fig. 1.**
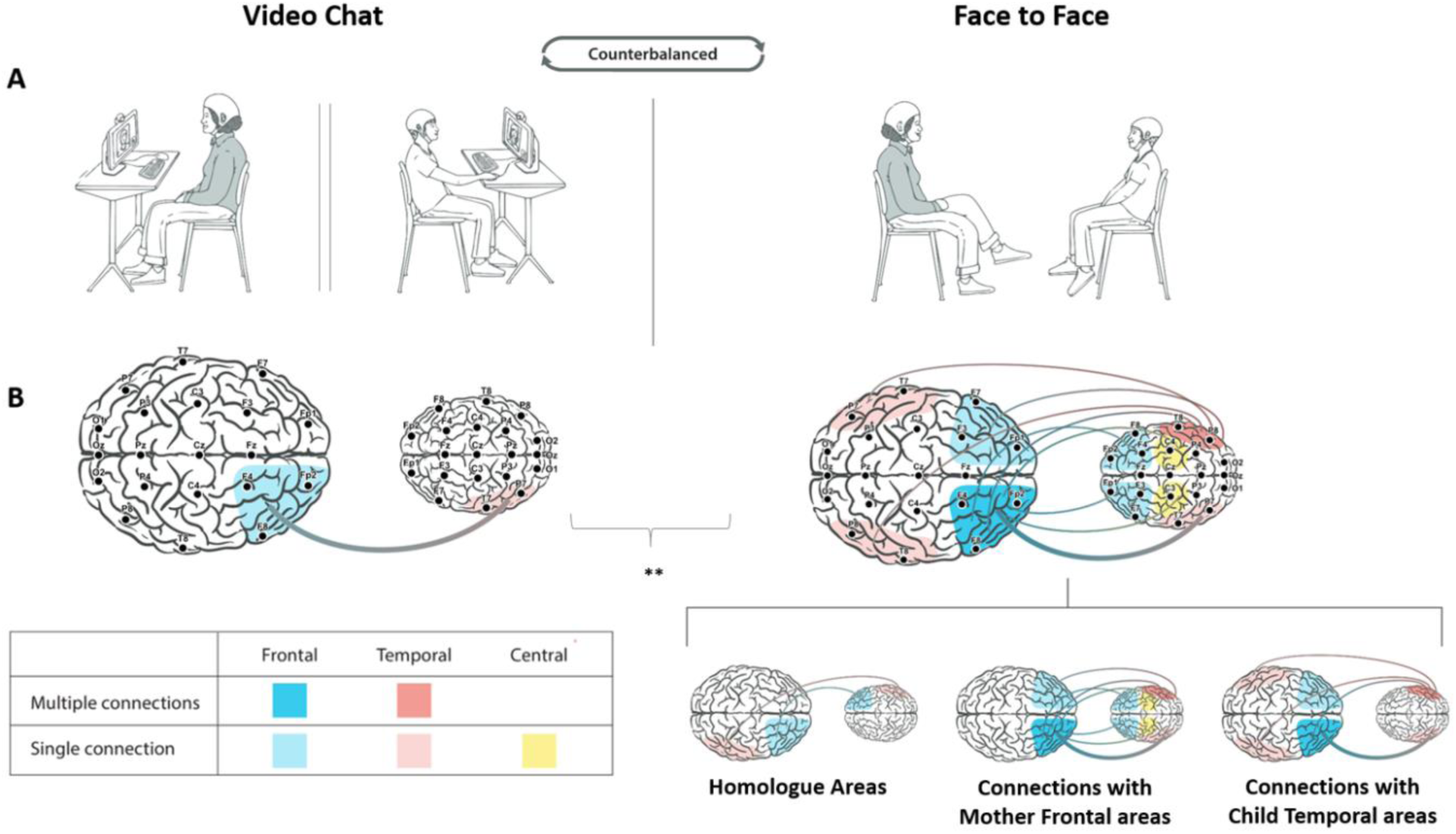
Experimental procedures and main findings. (A) Experiment paradigm. Neuroelectric activity of mother-child dyads was recorded simultaneously and continuously throughout the entire experiment (hypersacnning) during rest paradigm, face to face interaction, and while communicating via video chat form two separate locations (B) Main findings: Illustration of Mother-child inter-brain neural synchrony in both Face to face and Video chat paradigms. Inter-brain neural synchrony values were calculated for Beta frequency band (13.5 to 29.5 Hz) using weighted phase lag index (wPLI). 6 regions of interest were predefined and examined, each consisting of 3 electrodes. Region of interest were right and left Frontal, Central and Temporal areas. Connectivity scores were computed for the 6 regions of interest, resulting in 36 wPLI possible combinations between mother and child synchrony per condition. During the face to face interactions 9 inter-brain connections emerged between the mother and child, while in the video chat only one inter-brain connection was found. (*P <0.05 **P <0.01, ***P <0.001.)

Upon arrival, oral and written explanations of the procedures were given and participants signed informed consent. Following attachment to dual-EEG devices the mother and child sat 50 cm apart facing the wall and were instructed to look ahead and not interact for 2 minutes for the baseline (Rest) condition. Following the baseline paradigm, two counter-balanced positive-valance naturalistic interactions of three minutes each were recorded, during which mother and child were requested to either plan a fun day to spend together (option 1),plan a camping trip (option 2) or plan an amusement park visit (option 3). All options were counterbalanced across participants. In one interaction, the mother and child set and interacted face-to-face, and in the second they interacted via a video-chat from two separate rooms (Fig. 1A).

### 2.3 Dual neural and behavioral data acquisition

EEG activity of both mother and child was recorded simultaneously and continuously throughout the experiment. Data acquisition was performed using a 64-channels BrainAmp amplifier from Brain Products Company (Germany). The EEG system is composed of two BrainCap helmets including 32 electrodes each, arranged according to the international 10/20 system. The impedances were maintained below 10 kOhm and the ground electrode was placed on the AFz electrode. Both helmets were connected to the same amplifier to ensure millisecond-range synchrony between the EEG recording of the mother and child.

### 2.4 EEG Preprocessing

Preprocessing was conducted using Python 3.8, utilizing MNE software (v0.17.0). First, EEG data of each dyad was separated into two data files, one for the child and one for the mother, to enable separate preprocessing. Data was average referenced and a 1 to 50 Hz band-pass filter was applied on all data files, consistent with prior studies ((Djalovski et al., 2021; Endevelt-shapira et al., 2021), and, data was segmented into 1000 ms epochs with 500 ms overlap between epochs. Autoreject v0.1 (Jas et al., 2017) unsupervised algorithm with Bayesian optimization as the threshold method was utilized to remove trials containing transient jumps in isolated EEG channels and artifacts affecting groups of channels. Notably, while AutoReject specializes in excluding trials containing transient jumps in specific channels, systematic physiological artifacts that may affect multiple sensors, such as eye blinks or muscular movements is not optimally removed by this algorithm. Therefore, MNE’s implementations of Infomax and CORRMAP (Viola et al., 2009) were used to remove systematic physiological artifacts that affected the data. Independent components (IC) were manually selected for exclusion and served as templates for selecting and excluding similar components in all other participants across all the paradigms, so that the same IC templates were used across all conditions. Following preprocessing, we ascertained that the final number of epochs was similar across the three experimental conditions (see Supplementary Fig. S2).

### 2.5 Connectivity Analysis

Inter-brain synchrony was calculated using weighted phase lag index (wPLI), an inter-brain connectivity method that had been used in previous studies of naturalistic social interactions (Endevelt-shapira et al., 2021; Levy et al., 2017). The wPLI method reduces the probability of detecting “false positive” connectivity in case of a shared noise source, which may lead to false-positive hyper-connections resulting from similar sensory experiences for participants who are sharing the same settings and is considered an alternative method to PLV in naturalistic hyperscanning EEG studies (Burgess, 2013; Dikker et al., 2021). The wPLI method weighs each phase difference according to the magnitude of the lag so that phase differences around zero only marginally affect the calculation of the wPLI and was therefore suggested as an appropriate inter-brain connectivity method for assessing inter-brain connection during naturalistic social interactions. Importantly, while the wLPI was our main measure, we re-computed the data using the PLV to ascertain that all the cross-brain links found with the wLPI also show a significant linkage using the PLV method. Our results indicate that all face-to-face connections that were found using wPLI were indeed replicated using PLV. The results of this analysis are reported in Supplementary Table S1.

The dyad inter-brain neural connectivity values were calculated for the Beta frequency band (13.5-29.5 Hz). Consistent with prior research, we divided the EEG cap into pre-defined areas of interest based on the research hypotheses (Djalovski et al., 2021; Dumas et al., 2010), so that the EEG electrodes were grouped into predefined regions of interest (Djalovski et al., 2021; Kinreich et al., 2017), resulting in a total of 6 ROIs that were examined in this study, each consisting of 3 electrodes: right frontal (RF - Fp2, F4, F8), left frontal (LF - Fp1, F3, F7), right central (RC - FC2, CP2, C4), left central (LC - FC1, CP1, C3), right temporal (RT - T8, TP10, P8), and left temporal (LT - T7, TP9, P7). Overall, this resulted in 36 possible combinations of linkage between the mother’s and child’s ROIs.

Of the 70 dyads participating in the experiment, data files of 2 dyads were corrupted and discarded and 6 dyads did not share sufficient common epochs following AutoReject and IC rejection and connectivity could not be measured, resulting in a total of 62 dyads that were included in the analysis. Outlier values were excluded from analysis if values varied in more than 3 SDs from the mean of the delta connectivity calculated separately for each pair of ROIs for each analysis. No more than 3 participants were removed per analysis per comparison (face-to-face compare to rest analysis – exclusion average = 1.39 ± 0.64, video chat compared to rest analysis – exclusion average = 1.61 ± 0.6).

### 2.6 Behavioral Coding

To assess brain-behavior correlations, each paradigm was coded offline twice using two well-validated coding schemes: micro-coding and global rating using the Coding Interactive Behavior manual (CIB, Feldman, 1998).

The Interactive Behavior manual (CIB) is a well-validated rating system used for coding social interactions that has yielded over 200 publications across multiple cultures, age range, and pathological conditions (see Feldman, 2012, 2021 for review), including hyper-scanning research (Djalovski et al., 2021; Endevelt-shapira et al., 2021). The CIB yields 52 codes, each rated on 5-point scales that aggregate into theoretically-based constructs. Here, we used the *Child empathic social engagement* construct, which comprises the following CIB scales: Child gaze, child openness to parent, child involvement, child approach, child empathy, and child collaboration. *Child empathic social engagement* is a meaningful feature of the parent-child interaction at this age that differentiates healthy from pathological condition (Halevi et al., 2017), links with multiple hormones, including oxytocin, cortisol, and immune biomarkers (Yirmiya et al., 2020, 2018), and is individually stable from infancy to adolescence (Feldman, 2010). Each interactive context (face-to-face, video chat) was coded separately. Coding was conducted by trained coders who were blind to study hypotheses with inter-rater reliability for 20% of the interactions exceeding 90% on all codes (intra-class *r* = .93, range = .89-99).

The micro-coding utilized second-by-second coding of a previously validated coding scheme (Feldman and Eidelman, 2007, 2004) that has shown linkage with the brain basis of attachment in both parent (Atzil et al., 2014) and child (Pratt et al., 2018). Coding was conducted on a computerized system (Mangold Interact, Mangold International GmbH); gaze direction of each partner is coded separately and moments of shared gaze (gaze synchrony) are computed by the program.

### 2.7 Statistical Analysis

The neural connectivity statistical analysis was performed using *eelbrain*, an open source Python module for accessible statistical analysis of MEG and EEG data (v0.31.7, https://github.com/christianbrodbeck/eelbrain, DOI 10.5281/zenodo.598150). Connectivity was examined for the six ROIs for each dyad in each of the experimental condition.

For analyzing the differences between each of the experimental conditions, and to avoid multiple comparisons, we first used a non-parametric permutation test with mass-univariate ANOVA. We used a distribution derived permutation test since it is permuting the observed scores rather than assuming normal distribution. Thus, to analyze the data we first used a repeated-measures ANOVA to compute the F value for each of the ROI pairs, comparing between conditions within dyads. The same procedure was repeated in 1000 random permutations of the original data, shuffling condition labels within dyads to take into account the within-subject nature of the design. For each permutation, the largest F value was retained to form a nonparametric estimate of the distribution of the largest F value under the null hypothesis that condition labels are exchangeable. Last, a P value was computed for each ROI pair in the original F-map as the proportion of permutations that yielded a comparison with a larger F value than the comparison under question. Next, we used a more conservative method of a set of nonparametric Bonferroni corrected Wilcoxon signed-rank tests on all 36 possible Mother-Child ROI combinations in each condition. Only ROI pairs, which reached a P value of 0.05 or smaller following Bonferroni correction, are reported in the results section.

To further ensure the validity of our data, we conducted a set of two follow-up analysis: First, to validate that the observed differences in wPLI inter-brain connectivity are not related to changes in power (Marriott Haresign et al., 2022), beta power spectral density (PSD) was calculated for each of the mother and child ROIs and correlated with the inter-brain connectivity of the relevant ROI. PSD was calculated using MNE’s implementation of PSD using multitaper, and PSD scores were calculated for each electrode in each paradigm separately. Then, the power of each ROI was calculated as the average of the 3 relevant electrodes comprising each ROI. The power values of each ROI in each condition (face to face/ video chat) were then correlated with each of the wPLI connectivity values observed for the relevant ROI. Our results indicate that power did not correlate with any of the wPLI values in any of the experimental conditions in this study (see Supplementary Table S2). Therefore, the analysis revealed that power had no effect on the inter-brain connectivity values reported in our study, and the two separate measures were independent. In our next analysis we examined our data as compared to shuffled data: For each of the experimental paradigms (face to face/ video chat) we randomly shuffled the epochs of one member of each dyad 100 times and compared the original connectivity values of each of the significant inter-brain links reported in the results section to the connectivity values obtained from the shuffled data for that relevant ROI pair link. Once averaging the 100 iterations of the shuffled data, each of the inter-brain links in the original data was found to be significantly higher than the shuffled data (see Supplementary Fig. S1 for all significant links).

Finally, although we suggest wPLI is the more suited technique to measure inter-brain connectivity in natural settings, to further validate our main findings, we conducted a complimentary PLV connectivity analysis to further examine inter-brain synchrony between the 6 ROIs of the mother and child. PLV analysis results ensures that all significant connections found in the face-to face condition using wPLI were replicated using PLV. However, the video chat resulted in no significant linkages at all using PLV analysis (see Supplementary Table S1).

## 3. Results

### 3.1 Neural synchrony during live face-to-face interactions

We first compared neural synchrony during the live face-to-face interaction and baseline (rest condition) by using nonparametric permutation test with mass-univariate analysis of variance (ANOVA) based on one-way repeated-measures ANOVA designed to detect effects stemming from the face-to-face interaction as compared to Rest on wPLI scores. Results indicated a significant main effect of the face-to-face compared to baseline; (F(1,61) = 23.83, P = .001).

Following detection of the main effect, a more conservative method was used to calculate inter-brain connectivity between each ROI of the mother’s and child’s brains. Our following analysis included nonparametric Wilcoxon signed-rank tests to detect differences in wPLI connectivity between the face-to-face interaction and baseline. All results were Bonferroni-corrected to accommodate 36 comparisons. Significant (p < 0.05) inter-brain linkage was observed in 9 out of the possible 36 links (Fig. 1B, Fig 2A, Table 1, and Supplementary Fig. S3), which showed greater neural synchrony in the face-to-face interaction compared to the baseline and comprised four sub-groups: (a) *Homologous linkage between mother’s and child’s frontal and temporal regions,* (b) *inter-hemispheric same-region linkage of mother and child’s frontal and temporal regions,* (c) *linkage between mother’s frontal region and child’s temporal region*, and (d) *linkage between mother’s right frontal region and child’s central region*:

a. *Homologous linkage between mother’s and child’s frontal and temporal regions* – two homolog connectivity patterns were found: A right-temporal-right-temporal connectivity between mother and child (T = 450, z = 3.42, p_(Bonferroni corrected)_ = .022) and a right-frontal-right-frontal connectivity between mother and child (T = 364, z = 4.06, p_(Bonferroni corrected)_ = .002), both demonstrating linkage in the right hemisphere.
b. *Inter-hemispheric same-region linkage of mother and child’s frontal and temporal regions.* In addition to the homolog frontal and temporal connectivity, two cross-hemispheric links connected the mother’s and child frontal and temporal regions. Mother’s right frontal region linked with the child’s left frontal region (T = 389, z = 3.87, p_(Bonferroni corrected)_ = .004); and mother’s left temporal region linked with the child’s right temporal region (T = 483, z = 3.32 , p_(Bonferroni corrected)_ = .032), underscoring the tight connectivity of mother’s and child’s frontal and temporal regions.
c. *Linkage between mother’s frontal region and child’s temporal region*. Three inter-brain links connected mother’s frontal region and child’s temporal region. First, mother’s right frontal region linked with child’s right temporal region (T = 332, z = 4.29, p_(Bonferroni corrected)_ < .001); second, mother’s right frontal region linked with child’s left temporal region (T = 414, z = 3.69, p_(Bonferroni corrected)_ = .008); and third, mother’s left frontal region connected with child’s right temporal region (T = 428, z = 3.72, p_(Bonferroni corrected)_ = .007). These patterns highlight the tight cross-brain same and cross-hemisphere connectivity between the mother’s frontal region and the child’s temporal region.
d. *Linkage between mother’s right frontal region and child’s central region*. This included two links; between the mother’s right frontal region and the child’s left central region (T = 426, z = 3.60, p_(Bonferroni corrected)_ = .011), and between the mother’s right frontal region and the child’s right central region (T = 438, z = 3.51, p _Bonferroni (corrected)_ = .016) (Fig. 1B).

**Fig. 2.**
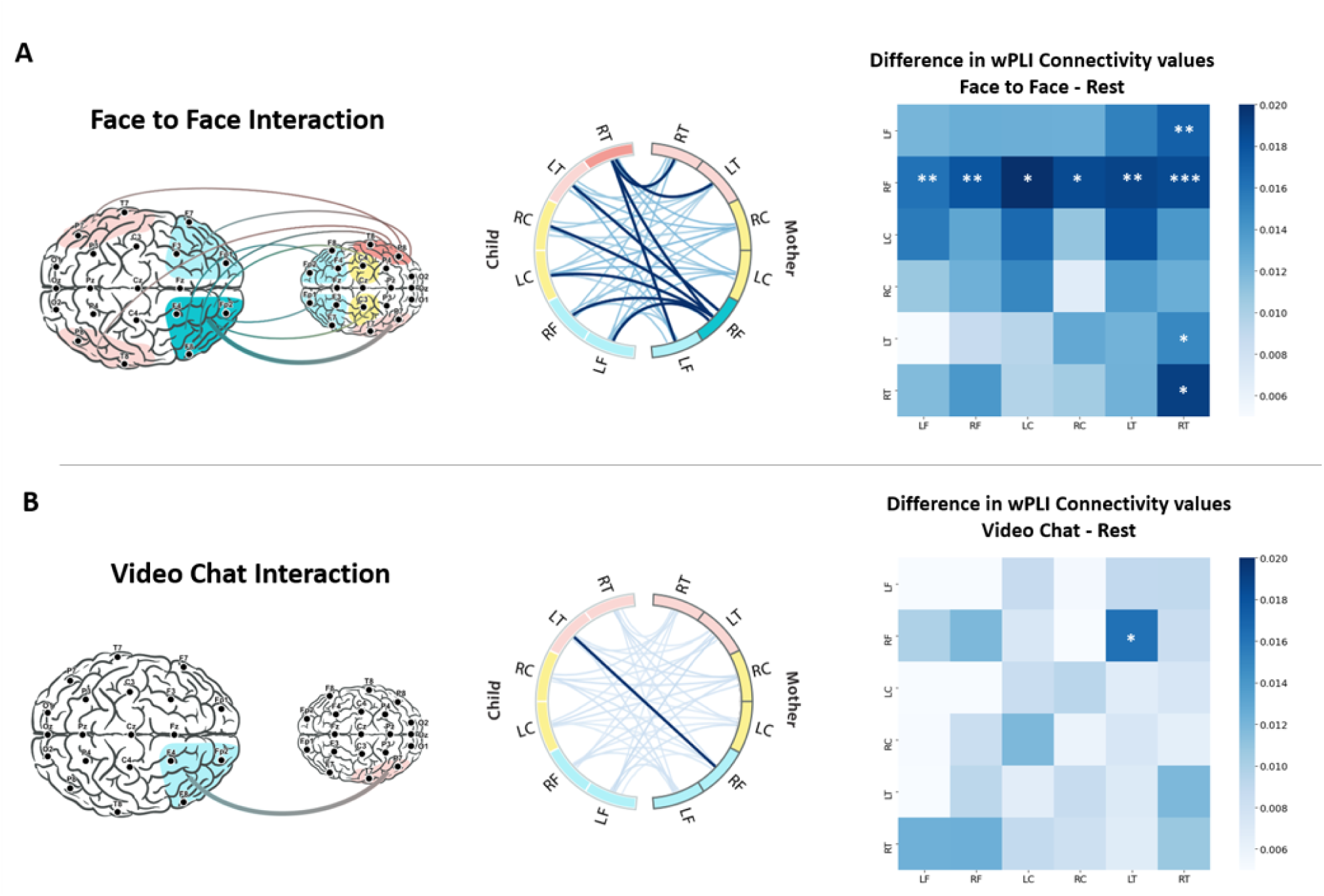
Visualization of significant Inter-brain Connections: Higher inter-brain synchrony was detected during face-to-face and Video chat interactions compared to rest condition, with 9 inter-brain connections found during face to face interaction, and a single connection in the video chat interaction. RT – right temporal, LT – left temporal, RC- right central, LC – left central, RF – right frontal, LF- left frontal. (A) Visualization of connectivity values (wPLI) during face to face interaction compared to rest. Circles represents mean connectivity values for the 36 possible combinations of region of interest in the mother and child brains. Within each circle, the significant links are marked in dark blue. Next, the difference in connectivity values across brain regions combinations between the face to face interaction and rest. The x axis represents the child’s brain region, while the y axis represents the mother’s brain regions. Darker squares represent comparisons with higher connectivity score differences between face to face and rest paradigms. Nonparametric permutation testing with mass-univariate ANOVA revealed significant main effect for face to face interaction compared to rest F(1,61) = 23.83 , P = .001). A conservative nonparametric Wilcoxon test was used to detect differences in wPLI connectivity measures, with all results Bonferroni-corrected to 36 comparison. 9 significant inter-brain connections were found post correction of the possible 36 combinations. The significant comparisons are marked. (B) Similar visualization of connectivity values (wPLI) during video chat interaction compared to rest condition. A single brain connection emerged between the Mother’s right frontal region and the child’s left temporal region following Bonferroni-corrections (F(1,61) = 17.11 , P = .001). (*P <0.05 **P <0.01, ***P <0.001.)

**Table 1:** Increased inter-brain neural synchrony following face to face interaction or Video chat interaction. Reported here are the significant comparisons following nonparametric Wilcoxon test to detect differences on wPLI connectivity measures between face to face or skype interactions and rest. All results were Bonferroni-corrected to 36 comparisons. Inter-brain neural synchrony was found in nine connections in the face to face paradigm, while a single inter-brain connection emerged in the video chat paradigm. *P <0.05 **P <0.01, ***P <0.001.

| Child ROI | Mother ROI | wPLI Face | wPLI Rest | p (FDR corrected) | p (Bonferroni corrected) | Sig (*) |
| --- | --- | --- | --- | --- | --- | --- |
| Face to Face Vs Rest |  |  |  |  |  |  |
| Right Frontal | Right Frontal | 0.118 (.031) | 0.100 (.026) | <.001 | 0.002 | ** |
| Right Temporal | Right Temporal | 0.117 (.035) | 0.098 (.022) | 0.003 | 0.022 | * |
| Left Frontal | Right Frontal | 0.115 (.030) | 0.099 (.021) | 0.001 | 0.004 | ** |
| Right Temporal | Left Temporal | 0.115 (.032) | 0.100 (.028) | 0.004 | 0.032 | * |
| Right Temporal | Right Frontal | 0.118 (.036) | 0.100 (.023) | <.001 | <.001 | *** |
| Left Temporal | Right Frontal | 0.117 (.033) | 0.098 (.021) | 0.002 | 0.008 | ** |
| Right Temporal | Left Frontal | 0.118 (.033) | 0.100 (.024) | 0.002 | 0.007 | ** |
| Left Central | Right Frontal | 0.12 (.040) | 0.098 (.022) | 0.002 | 0.011 | * |
| Right Central | Right Frontal | 0.12 (.038) | 0.101 (.022) | 0.002 | 0.016 | * |
| Video Chat Vs Rest |  |  |  |  |  |  |
| Left Temporal | Right Frontal | 0.113 (.031) | 0.100 (.029) | 0.02 | 0.02 | * |

As seen, a wide net of connections links mother and child’s brains during live social interactions. Particularly salient in these connectivity patterns are the mother’s right frontal and the child’s right temporal regions. Mother’s *right frontal region* connected with *each of the six child’s ROIs measured here*; the child’s right and left frontal, right and left temporal, and right and left central regions. Similarly, the child’s *right temporal region* was multiply connected; with the mother’s right and left frontal regions and right and left temporal regions. Notably, the mother’s frontal and child’s temporal areas showed not only same-hemisphere linkage but also cross-hemispheric connectivity, highlighting these two areas in mother and child as densely inter-connected during live social exchanges.

### 3.2 Neural synchrony during technologically-assisted communication

The next analysis compared inter-brain synchrony during the video chat compared to the baseline condition using the same analysis; nonparametric permutation test with mass-univariate analysis of variance (ANOVA) based on one-way repeated-measures ANOVA design to detect effects associated with the video chat free interaction compared to the rest condition on wPLI scores. Results indicated a significant overall main effect for the video chat compared to the rest (F(1,61) = 17.11, P = .001). A follow-up nonparametric Bonferroni-corrected Wilcoxon tests to detect differences on wPLI connectivity measures corrected for 36 comparisons was conducted, similar to the face-to-face condition. Following Bonferroni correction, only a single significant inter-brain link emerged of the possible 36 links; between the mother’s right frontal region and the child’s left temporal region (T = 446, z = 3.45, p_(Bonferroni corrected)_ = .02) (Table 1, Fig. 2B). Overall, our findings demonstrate the marked decrease in brain-to-brain connectivity when interactions are moderated by technology. The single inter-brain link found during the video chat further underscores the mother-frontal-child-temporal link as central for supporting mother-child social interactions during both during live or remote communication.

### 3.3 Comparing inter-brain connectivity during live versus technologically-assisted communication

Z-test for Two Proportions was used to examine differences between the statically significant inter-brain connections during the face-to-face and video chat. A significant difference was found between the overall inter-brain connections during the live versus remote interaction (Z = 2.27, *p* = .006), supporting our main hypothesis of a robust decrease in inter-brain synchrony when interactions are mediated by technology.

### 3.4 Brain-behavior coupling

Consistent with previous studies indicating brain-behavior coupling with homolog connectivity patterns (Djalovski et al., 2021; Kinreich et al., 2017), we examined associations between gaze synchrony and empathic social engagement with the temporal-temporal and frontal-frontal links.

*Gaze synchrony links with live temporal-temporal synchrony* – Moments of shared gaze between mother and child – gaze synchrony – correlated with mother-child temporal-temporal link during the live interaction, r = 0.28, P = 0.032 (Fig. 3A); more gaze synchrony correlated with greater right temporal connectivity. Gaze synchrony was unrelated to temporal-temporal synchrony in the video chat (r = 0.15, P > 0.25); still, Fisher’s Z transformation examining the difference between these two correlations was not significant (Z = 0.79, P = 0.21). Finally, Mother-child frontal-frontal connectivity was unrelated to gaze synchrony in both the live (r = 0.23, P = 0.083(, or remote (r = 0.23, P = 0.097) communication conditions.

**Fig. 3.**
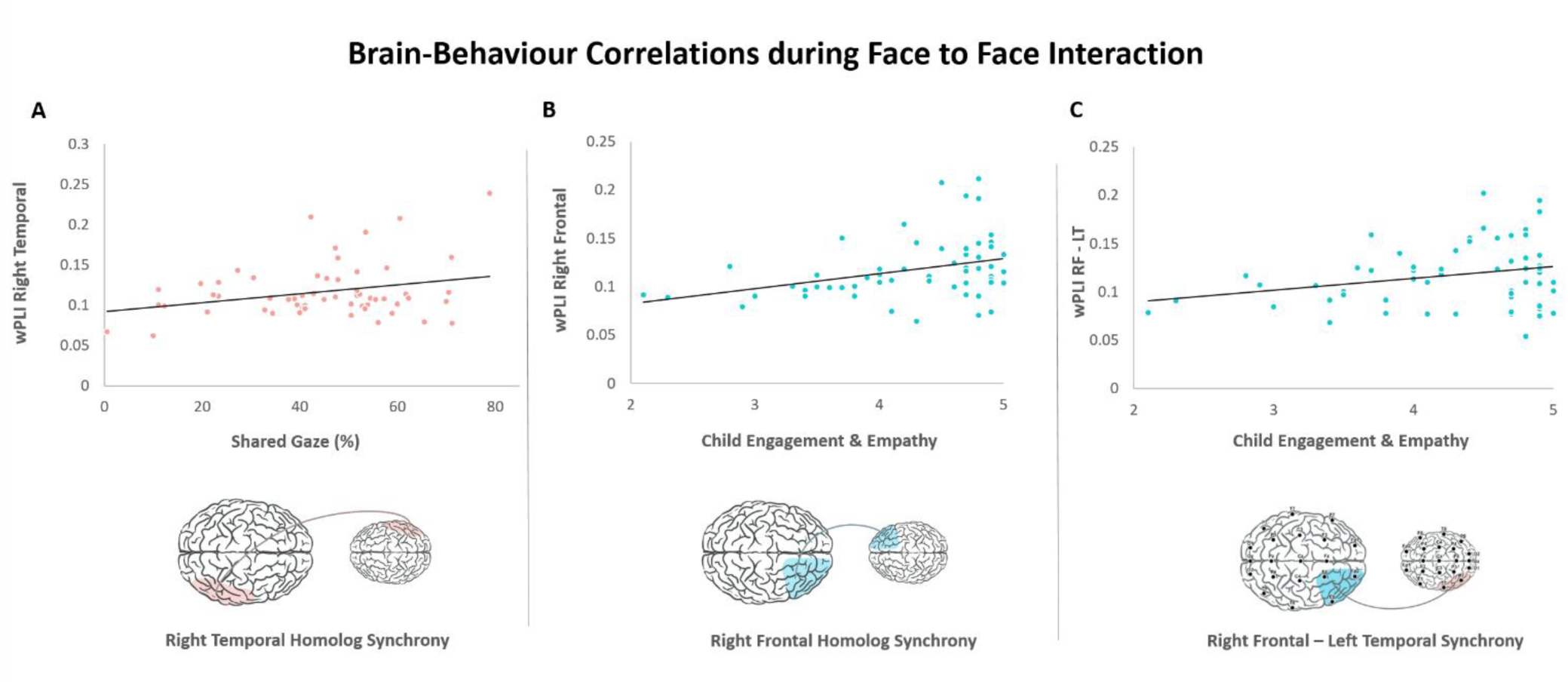
Visualization of brain-behaviour correlations in the Face to Face Paradigm. (A) Visualization of mother-child dyads shared gaze during interactions correlation with wPLI values of inter-brain synchrony. Neural synchrony was highly correlated with synchronous gaze shared between the mother and child in the right temporal area (r = 0.28, P = 0.032). (B + C) Visualization of the CIB codes of child Engagements and Empathy correlations with wPLI values of inter-brain neural synchrony. The extent of child engagement and empathy towards the mother affected synchrony in the homolog frontal right areas of the mother and child (r = 0.35, p = 0.007), and in the frontal right area of the mother with the temporal left area of the child (r = 0.26, P = 0.043).

C*hild empathic social engagement links with live frontal-frontal synchrony -* child empathic social engagement correlated with mother-child frontal-frontal connectivity, r = 0.35, *P* = 0.007 (Fig. 3B), but not during the video chat interaction, r = 0.06, P > 0.25. Fisher’s Z transformation test showed significant difference between the magnitude of the correlations (Z = 1.7, *P* = 0.045).

Following the a-priori assessment of brain-behavior coupling in homolog connectivity patterns, we also used a post-hoc exploratory analysis to examine the association between *Child empathic engagement* and the single inter-brain link that was significant during both the live and video chat interactions. Results showed that the mother-right-frontal-child-left-temporal link correlated with *child empathic engagement* during the live interactions; r = 0.26, P = 0.043 (Fig. 3C), but not during the video chat, r = -0.03, P > 0.25. Following Fisher’s Z transformation, the two correlations showed marginally significant difference (Z = 1.62, *P* = 0.052). As hypothesized, while the micro- and macro behavioral constructs correlated with neural coordination in the live face-to-face interaction, brain-behavior correlations during the technologically-assisted communication were not significant

## 4 Discussion

The COVID-19 pandemic has accelerated an already ongoing revolution of proportion and consequences unbeknown to our species, making remote communication a topic of urgent public concern. Our study is the first to tackle this issue from a two-brain perspective that focuses on inter-brain synchrony, a key mechanism sustaining human social connections (Feldman, 2021, 2020). The findings clearly demonstrate the price we pay for technology. As shown, during mother-adolescent live face-to-face interaction a wide net of connections unfolds between the two brains, including (a) homolog same-region-same-hemisphere links; (b) same-region-different-hemisphere links; and (c) multi-region patterns of connectivity. Particularly salient were links between the mother’s and the child’s frontal and temporal regions, which were inter-connected in nearly every possible way and rode on beta rhythms. In contrast to the 9 significant links working in tandem during the live interaction, only 1 significant link connected the two brains during remote communication: between the mother’s right frontal region and the child’s left temporal region. Remote interaction, therefore, eliminates the rich right-to-right brain linkage repeatedly found in naturalistic cross-brain studies (Cui et al., 2012; Kruppa et al., 2021; Pan et al., 2017; Reindl et al., 2018) and theorized to transmit the partners’ non-verbal social cues and affective states (Borod et al., 1998). Furthermore, while inter-brain synchrony during the live interaction was embedded in social behavior, no significant brain-behavior correlations emerged during the video chat, resulting in a thinned bio-behavioral experience. Overall, our findings suggest that the substantial gains to social development, empathic abilities, and brain maturation afforded by face-to-face interactions may not translate to technological encounters, opening a much needed discussion on the price of technological communication. Moreover, we need to understand whether the "zoom fatigue" we feel after several moments of remote communication, possibly due to the immense load placed on a single channel of inter-brain linkage, impairs the development of children’s sustained attention and recognition of affective cues; what are the biological mechanisms implicated in "co-presence" between two humans; and what may be the long-term effects of significant reduction in live social interactions on maturation of the social brain, particularly during key developmental transitions such as the transition to adolescence.

During the mother-child face-to-face interaction, two homolog inter-brain connections were found; between mother and child’s (1) right frontal regions and (2) right temporal regions. The frontal linkage is consistent with much prior research. A recent rodent study demonstrated the causal involvement of frontal inter-brain synchrony in sociality; when frontal neurons were activated synchronously the animals were socially engaged but when the same neurons activated asynchronously, they lost social interest (Yang et al., 2022). Hyperscanning studies have similarly shown that right frontal-frontal synchrony sustains human affiliation, including parent-child (Kruppa et al., 2021; Reindl et al., 2018) and romantic attachment (Pan et al., 2017); frontal-frontal synchrony decreases when mothers are stressed (Azhari et al., 2019); and frontal-frontal synchrony links with perceived similarity between partners (Hu et al., 2017) and sense of effective communication (Stephens et al., 2010). Here we found that the mother’s frontal region linked with every single region of the child’s brain measured; child’s right and left frontal, right and left central, and right and left temporal areas (Fig. 1B), suggesting a unique role for the mother’s right frontal area in sustaining inter-brain synchrony. The mother’s right frontal region may be involved in monitoring the interaction and dynamically adjusting its features to ensure rich inter-brain coupling at multiple levels of the child’s neural processing. The frontal cortex is implicated in higher-order social functions, including social cognition, mental state knowledge, and social decision-making (Amodio and Frith, 2006; Rilling and Sanfey, 2011), abilities that are known to develop in the context of maternal care (Monroy et al., 2010). The dense cross-brain linkage emanating from the mother’s right frontal cortex accords with the well-known mechanism of "external regulation" (Hofer, 1995), the process by which the mature maternal brain molds the child’s immature brain and tunes it to social life through inter-brain mechanisms embedded within coordinated social behavior (Feldman, 2021, 2015).

The right temporal-temporal link is similarly consistent with previous research during interactions between attachment partners, suggesting its role in the formation of affiliative bonds (Djalovski et al., 2021; Kinreich et al., 2017). The right temporal region is involved in empathy, embodiment, and mentalization and underpins the capacity to understand others’ goals and create shared intentionality during social moments (Frith and Frith, 2001). Increased beta activations were found in right temporal regions when children observe their own mother-child videos (Pratt et al., 2018), when mothers are exposed to infant-related emotional stimuli (Hernández-González et al., 2016), and when romantic partners engage in empathic dialogue (Djalovski et al., 2021) and it has been suggested that temporal beta serves as a neural marker of attachment (Hernández-González et al., 2016). During adolescence, the social brain undergoes profound reorganization in both the PFC and the posterior superior temporal sulcus (Blakemore, 2008). We found that during a period of such rapid maturation of these areas, moments of naturalistic mother-adolescent social interaction trigger not only a homolog right-brain linkage of these areas, but also a dense inter-connection between mother and child’s right and left frontal and temporal regions. The fronto-temporal network underpins key socio-cognitive functions (Frith and Frith, 2001; Hastings et al., 2013), and hyperscanning studies indicated frontal-temporal neural synchrony during social exchanges (Pérez et al., 2017; Tang et al., 2015; Zhang et al., 2017). This suggests that mothers utilize inter-brain mechanisms to support maturation of the social brain during its sensitive periods of development in stage-specific ways that target the specific areas that undergo rapid development, findings that lend further support to the perspective that inter-brain synchrony is a mechanism by which mature brains regulate immature brains to social living (Feldman, 2020, 2016).

All inter-brain links found here implicated beta rhythms. Inter-brain processes are sustained by neural oscillations, a pervasive component of neuronal activity that underpins the dynamic organization of neural functions (Donner and Siegel, 2011), and their temporal consistency builds a model of self and partner’s behavior that can guide neuronal activity toward a smooth interpersonal exchange (Seth and Friston, 2016). Beta oscillations are involved in post-synaptic gains in neuronal sensitivity that modify predictions and determine information flow towards higher-order targets (Bressler and Richter, 2015; Friston et al., 2015). Beta rhythms are involved in complex social functions, such as empathy (Levy et al., 2018) and attachment (Pratt et al., 2018), and underpin key functions that enable cross-brain communication, including active information processing (Donner and Siegel, 2011), mentalization (Soto-Icaza et al., 2019), predicting others’ actions (Koelewijn et al., 2008), perception and integration of sensory information (Hipp et al., 2011), and the constant adaptations and updating of predictions (Sedley et al., 2016). During social interactions, these beta-modulated functions enable the rapid adaptation and mutual entrainment that are required for inter-brain coordination (Hasson and Frith, 2016).

Hyperscanning studies revlealed cross-brain synchrony of beta rhythms across multiple tasks, such as response to positive social gestures (Balconi and Fronda, 2020), compassion during third-party punishment (Ciaramidaro et al., 2018), and leader-follower cooperation (Yun et al., 2012). Inter-brain beta synchrony has been found during synchronized movements and the increase in inter-brain beta during episodes of coordinated movement was interpreted as representing top-down modulations in social interactions that derive from joint action, social attention, and imitation (Dumas et al., 2010). Enhanced beta-band synchronization emerged during cooperation, as compared to competition paradigms when partners are co-present (Sinha et al., 2016). Furthermore, consistent with the current results, inter-brain beta during cooperation was found in both frontal and right-temporo-parietal areas (Sciaraffa et al., 2021) and was suggested to derive from the active thinking, joint focus, and metallizing processes that are triggered by coordination dynamics. Finally, interpersonal factors, such as trait empathy, engagement, and social behavior of joint engagement and eye contact were found to predict inter-brain beta during real-world face-to-face interactions (Dikker et al., 2021).

Consistent with the bio-behavioral synchrony model (Feldman, 2016, 2015, 2012), inter-brain synchrony was associated with behavioral coordination and engagement during the live, but not the remote interaction. The partners’ right temporal-temporal link increased as a function of their shared gaze, which serves as a critical social cue and is the main nonverbal channel of communication. Extant inter-brain literature demonstrated the effects of shared gaze on facilitating cross-brain synchrony (Endevelt-shapira et al., 2021; Hirsch et al., 2017; Kinreich et al., 2017; Koike et al., 2019; Leong et al., 2017; Piazza et al., 2020). Shared gaze has been suggested to enhance neural coordination by supporting the ability to communicate social signals, predict ongoing intent, identify partner’s affective state, and execute a joint goal (Schilbach et al., 2013; Tang et al., 2015). Hperscanning EEG studies demonstrated both frontal and temporal connectivity during face-to-face interaction involving shared gaze (Hirsch et al., 2017; Markova et al., 2019; Noah et al., 2020), and naturalistic interactions triggered right temporal-temporal synchrony that were embedded in moments of gaze synchrony (Kinreich et al., 2017). Similarly, face-to-face interactions containing shared gaze triggered more neural synchrony than a pre-filmed video of interacting faces (Noah et al., 2020). In mother-infant dyads, moments of shared gaze, co-vocalizations, and joint affect facilitate inter-brain synchrony (Markova et al., 2019); neural synchrony increases during face-to-face compared to back-to-back (Endevelt-shapira et al., 2021); and shared gaze triggers greater neural coupling compared to averted or no gaze (Leong et al., 2017; Piazza et al., 2020). Our findings add three important points to this literature. First, results pinpoint the association between shared gaze and right temporal-temporal synchrony; second, they implicate beta rhythms in mother-adolescent bio-behavioral synchrony, and third, although the degree of shared gaze was comparable between the live and remote communication, only during the live interaction did this fundamental, first to mature social cue of shared gaze was significantly correlated with inter-brain coupling.

While inter-brain synchrony during infancy and early childhood mainly link with the mother’s behavior, such as touch, gaze, or vocalizations (Leong et al., 2017; Markova et al., 2019), results highlight the role of children’s social behavior for neural coupling at the transition to adolescence, possibly representing a shift in the relational dynamics as children enter the second decade of life. Our data reveal that children’s involvement in the dialogue, empathy, collaboration, and social motivation, reflected in the *empathic social engagement* construct, facilitated mother-child right frontal-frontal synchrony during live interactions. This shift may reflect developmental processes underpinning social brain maturation during the adolescent transition and the growing capacities for empathy associated with this shift. The expressions of child empathy and collaboration, higher-order abilities known to sustain inter-brain synchrony, were comparable during the live and remote communication, still, these social behaviors connected with greater frontal-frontal and frontal-temporal synchrony only in the live interaction and showed no significant correlations with neural coupling in the remote communication. As such, our results suggest that live social interactions provide the evolutionary-typical context for the maturation of neural coupling, findings that raise concerns about the rates of youth involvement in technologically-assisted communication and the potential risk this poses to the development empathy and collaboration.

Although adolescents are accustomed to technological communication (Anderson and Jiang, 2018), remote learning is problematic; US junior high school students are estimated to acquire 37-65% of the yearly skills during remote learning (Kuhfeld et al., 2020). The exact reasons for such impoverished learning are not fully clear and factors such as delay in social feedback, difficulties in maintaining attention, faceless participants, or mute microphones were suggested to aggravate zoom fatigue (Peper et al., 2021). Here we show that even under the best circumstances – partners are familiar with each other’s signals, technology is fully adjusted, and the topic is a relaxed conversation – inter-brain coordination is significantly severed during remote contact. Our findings may suggest that one reason for the zoom fatigue continuously reported during the COVID-19 pandemic is that remote communication traffics through a single inter-brain link, while the brain is used to a dense net of connections that transmit information at various levels across the neuroaxis: sensory, motor, linguistic, affective, and shared meaning, with each level possibly riding on a distinct cross-brain link. Furthermore, it appears that whereas during live interactions the brain can integrate information from social behavior and empathic resonance, social behaviors may be less helpful for the creation of shared brain states when partners interact remotely, rendering the creation of shared meaning a more effortful task.

We are only beginning to understand how social technology impacts the human social brain and this topic is in urgent need of further research. We need to understand the cross-brain consequences of technological communication at different stages of child development and with different familiar and unfamiliar partners. As technologically-assisted communication is assuming an increasing portion of our social life, we must address the broader implications of this change; how remote communication impacts parenting, falling in love, couple relationships, social communities, self-identity, and resilience. We must learn to quantify the amount of technologically-assisted communication that may tilt the developing brain to less favorable outcomes at each stage of development. Finally, a key goal for future technology and research is to test whether there are components of the human biological co-presence that can be adapted to screen-mediated interaction. Technology offers a wealth of possibilities and, with it, the option to alter what it means to be a social human. More empirical knowledge may help address these questions with wisdom and foresight for the future development of tomorrow’s citizens.

## Acknowledgements

The study was supported by the Simms/Mann Foundation Chair to Ruth Feldman and by the Bezos Family Foundation.

## Notes

### Competing Interest Statement

The authors have declared no competing interest.

