## Supplementary material for "Technologically-Assisted Communication Attenuates Inter-Brain Synchrony"

**Table S1 – PLV analysis**

| Sig (*) | p (FDR corrected) | PLV Rest | PLV Face-to-Face | Mother ROI | Child ROI |
| --- | --- | --- | --- | --- | --- |
| * | .0120 | 0.067 *(.013)* | 0.076 *(.022)* | Right Frontal | Right Frontal |
| ** | 0.008 | 0.067 *(.012)* | 0.076 *(.023)* | Right Temporal | Right Temporal |
| * | 0.012 | 0.067 *(.012)* | 0.076 *(.024)* | Right Frontal | Left Frontal |
| * | 0.011 | 0.066 *(.013)* | 0.075 *(.024)* | Left Temporal | Right Temporal |
| * | 0.039 | 0.068 *(.017)* | 0.076 *(.025)* | Right Frontal | Right Temporal |
| ** | 0.006 | 0.067 *(.013)* | 0.076 *(.023)* | Right Frontal | Left Temporal |
| ** | 0.006 | 0.066 *(.012)* | 0.077 *(.024)* | Left Frontal | Right Temporal |
| ** | 0.008 | 0.067 *(.014)* | 0.078 *(.024)* | Right Frontal | Left Central |
| ** | 0.008 | 0.068 *(.016)* | 0.076 *(.023)* | Right Frontal | Right Central |
| Sig (*) | p (FDR corrected) | PLV Rest | PLV video-chat | Mother Area | Child Area |
| n.s. | 0.026 | 0.07 *(.023)* | 0.071 *(.018)* | Right Frontal | Left Temporal |

**Table S1. Phase locking value (PLV) analysis revealed that all significant links found using wPLI were replicated and found significant using PLV as well in the Face to Face compared to baseline analysis.** To further validate our main findings, we conducted a PLV connectivity analysis to examined inter-brain synchrony between the 6 ROIs of the mother and child. Notably, the video chat resulted in no significant linkages at all using PLV analysis. (*P <0.05 **P <0.01, ***P <0.001.)

**Table S2 – Correlations between Power and Connectivity across ROIs and Paradigms**

**Correlations between Power and Connectivity – Face to Face Paradigm – Mother's ROIs**

| **p value (uncorrected)** | **t value** | **r'** | **wPLI Connectivity** | |  | **Power** | **ROI** |
| --- | --- | --- | --- | --- | --- | --- | --- |
| 0.067 | -1.867 | -0.234 | 0.116 |  | RT (mother) + RT (child) | -121.709 | RT (mother) |
| 0.794 | -0.262 | -0.034 | 0.113 |  | RT (mother) + LT (child) |  |  |
| 0.757 | 0.311 | 0.040 | 0.114 |  | RT (mother) + RF (child) |  |  |
| 0.382 | -0.881 | -0.113 | 0.114 |  | RT (mother) + LF (child) |  |  |
| 0.395 | -0.858 | -0.110 | 0.113 |  | RT (mother) + RC (child) |  |  |
| 0.178 | -1.361 | -0.173 | 0.111 |  | RT (mother) + LC (child) |  |  |
| 0.389 | -0.867 | -0.111 | 0.114 |  | LT (mother) + RT (child) | -120.550 | LT (mother) |
| 0.212 | -1.261 | -0.161 | 0.112 |  | LT (mother) + LT (child) |  |  |
| 0.617 | -0.502 | -0.065 | 0.110 |  | LT (mother) + RF (child) |  |  |
| 0.420 | -0.811 | -0.104 | 0.109 |  | LT (mother) + LF (child) |  |  |
| 0.466 | -0.734 | -0.094 | 0.111 |  | LT (mother) + RC (child) |  |  |
| 0.592 | -0.540 | -0.069 | 0.111 |  | LT (mother) + LC (child) |  |  |
| 0.171 | -1.386 | -0.176 | 0.117 |  | RF (mother) + RT (child) | -124.958 | RF (mother) |
| 0.277 | -1.096 | -0.140 | 0.116 |  | RF (mother) + LT (child) |  |  |
| 0.341 | -0.960 | -0.123 | 0.117 |  | RF (mother) + RF (child) |  |  |
| 0.120 | -1.576 | -0.199 | 0.117 |  | RF (mother) + LF (child) |  |  |
| 0.428 | -0.797 | -0.102 | 0.118 |  | RF (mother) + RC (child) |  |  |
| 0.280 | -1.089 | -0.139 | 0.119 |  | RF (mother) + LC (child) |  |  |
| 0.977 | 0.029 | 0.004 | 0.117 |  | LF (mother) + RT (child) | -124.748 | LF (mother) |
| 0.705 | -0.380 | -0.049 | 0.118 |  | LF (mother) + LT (child) |  |  |
| 0.778 | 0.284 | 0.037 | 0.114 |  | LF (mother) + RF (child) |  |  |
| 0.517 | -0.651 | -0.084 | 0.114 |  | LF (mother) + LF (child) |  |  |
| 0.812 | -0.239 | -0.031 | 0.115 |  | LF (mother) + RC (child) |  |  |
| 0.739 | -0.334 | -0.043 | 0.110 |  | LF (mother) + LC (child) |  |  |
| 0.277 | 1.098 | 0.140 | 0.116 |  | RC (mother) + RT (child) | -127.721 | RC (mother) |
| 0.531 | -0.630 | -0.081 | 0.116 |  | RC (mother) + LT (child) |  |  |
| 0.641 | -0.468 | -0.060 | 0.113 |  | RC (mother) + RF (child) |  |  |
| 0.547 | -0.605 | -0.078 | 0.114 |  | RC (mother) + LF (child) |  |  |
| 0.823 | -0.224 | -0.029 | 0.111 |  | RC (mother) + RC (child) |  |  |
| 0.520 | -0.646 | -0.083 | 0.118 |  | RC (mother) + LC (child) |  |  |
| 0.854 | 0.184 | 0.024 | 0.117 |  | LC (mother) + RT (child) | -128.456 | LC (mother) |
| 0.860 | 0.177 | 0.023 | 0.113 |  | LC (mother) + LT (child) |  |  |
| 0.917 | -0.105 | -0.014 | 0.113 |  | LC (mother) + RF (child) |  |  |
| 0.508 | -0.666 | -0.086 | 0.116 |  | LC (mother) + LF (child) |  |  |
| 0.798 | -0.256 | -0.033 | 0.111 |  | LC (mother) + RC (child) |  |  |
| 0.197 | -1.303 | -0.166 | 0.117 |  | LC (mother) + LC (child) |  |  |

**Correlations between Power and Connectivity – Face to Face paradigm – Child's ROIs**

| **p value (uncorrected)** | | **t value** | | **r'** | | **wPLI Connectivity** | |  | | **Power** | | **ROI** |
| --- | --- | --- | --- | --- | --- | --- | --- | --- | --- | --- | --- | --- |
| 0.863 | | -0.174 | | -0.022 | | 0.116 | | RT (mother) + RT (child) | | -119.347 | | RT (child) |
| 0.866 | | 0.169 | | 0.022 | | 0.114 | | LT (mother) + RT (child) | |  | |  |
| 0.863 | | 0.174 | | 0.022 | | 0.117 | | RF (mother) + RT (child) | |  | |  |
| 0.922 | | 0.098 | | 0.013 | | 0.117 | | LF (mother) + RT (child) | |  | |  |
| 0.645 | | 0.463 | | 0.060 | | 0.116 | | RC (mother) + RT (child) | |  | |  |
| 0.692 | | 0.398 | | 0.051 | | 0.117 | | LC (mother) + RT (child) | |  | |  |
| 0.400 | | 0.848 | | 0.109 | | 0.113 | | RT (mother) + LT (child) | | -120.074 | | LT (child) |
| 0.765 | | -0.301 | | -0.039 | | 0.112 | | LT (mother) + LT (child) | |  | |  |
| 0.126 | | 1.550 | | 0.196 | | 0.116 | | RF (mother) + LT (child) | |  | |  |
| 0.433 | | 0.789 | | 0.101 | | 0.118 | | LF (mother) + LT (child) | |  | |  |
| 0.400 | | 0.848 | | 0.109 | | 0.116 | | RC (mother) + LT (child) | |  | |  |
| 0.136 | | 1.509 | | 0.191 | | 0.113 | | LC (mother) + LT (child) | |  | |  |
| 0.224 | | 1.227 | | 0.157 | | 0.114 | | RT (mother) + RF (child) | | -123.000 | | RF (child) |
| 0.007* | | 2.783 | | 0.338 | | 0.110 | | LT (mother) + RF (child) | |  | |  |
| 0.307 | | 1.029 | | 0.132 | | 0.117 | | RF (mother) + RF (child) | |  | |  |
| 0.113 | | 1.610 | | 0.203 | | 0.114 | | LF (mother) + RF (child) | |  | |  |
| 0.230 | | 1.213 | | 0.155 | | 0.113 | | RC (mother) + RF (child) | |  | |  |
| 0.141 | | 1.494 | | 0.189 | | 0.113 | | LC (mother) + RF (child) | |  | |  |
| 0.483 | | 0.706 | | 0.091 | | 0.114 | | RT (mother) + LF (child) | | -123.403 | | LF (child) |
| 0.043* | | 2.070 | | 0.258 | | 0.109 | | LT (mother) + LF (child) | |  | |  |
| 0.661 | | 0.440 | | 0.057 | | 0.117 | | RF (mother) + LF (child) | |  | |  |
| 0.129 | | 1.537 | | 0.195 | | 0.114 | | LF (mother) + LF (child) | |  | |  |
| 0.043* | | 2.064 | | 0.257 | | 0.114 | | RC (mother) + LF (child) | |  | |  |
| 0.299 | | 1.047 | | 0.134 | | 0.116 | | LC (mother) + LF (child) | |  | |  |
| 0.863 | | 0.174 | | 0.022 | | 0.113 | | RT (mother) + RC (child) | | -125.887 | | RC (child) |
| 0.342 | | 0.957 | | 0.123 | | 0.111 | | LT (mother) + RC (child) | |  | |  |
| 0.823 | | -0.225 | | -0.029 | | 0.118 | | RF (mother) + RC (child) | |  | |  |
| 0.421 | | 0.811 | | 0.104 | | 0.115 | | LF (mother) + RC (child) | |  | |  |
| 0.625 | | 0.491 | | 0.063 | | 0.111 | | RC (mother) + RC (child) | |  | |  |
| 0.691 | | 0.399 | | 0.051 | | 0.111 | | LC (mother) + RC (child) | |  | |  |
| 0.534 | | 0.625 | | 0.080 | | 0.111 | | RT (mother) + LC (child) | | -126.411 | | LC (child) |
| 0.775 | | 0.287 | | 0.037 | | 0.111 | | LT (mother) + LC (child) | |  | |  |
| 0.611 | | 0.511 | | 0.066 | | 0.119 | | RF (mother) + LC (child) | |  | |  |
| 0.724 | | -0.355 | | -0.046 | | 0.110 | | LF (mother) + LC (child) | |  | |  |
| 0.128 | | 1.543 | | 0.195 | | 0.118 | | RC (mother) + LC (child) | |  | |  |
| 0.956 | | 0.055 | | 0.007 | | 0.117 | | LC (mother) + LC (child) | |  | |  |

**Correlations between Power and Connectivity – Video chat Paradigm – Mother's ROIs**

| **p value (uncorrected)** | **t value** | **r'** | **wPLI Connectivity** |  | **Power** | **ROI** |
| --- | --- | --- | --- | --- | --- | --- |
| 0.147 | -1.469 | -0.186 | 0.116 | RT (mother) + RT (child) | -121.344 | RT (mother) |
| 0.578 | -0.559 | -0.072 | 0.120 | RT (mother) + LT (child) |  |  |
| 0.173 | -1.379 | -0.175 | 0.116 | RT (mother) + RF (child) |  |  |
| 0.216 | -1.250 | -0.159 | 0.116 | RT (mother) + LF (child) |  |  |
| 0.172 | -1.381 | -0.176 | 0.118 | RT (mother) + RC (child) |  |  |
| 0.380 | -0.885 | -0.114 | 0.114 | RT (mother) + LC (child) |  |  |
| 0.805 | 0.247 | 0.032 | 0.120 | LT (mother) + RT (child) | -120.760 | LT (mother) |
| 0.927 | 0.091 | 0.012 | 0.116 | LT (mother) + LT (child) |  |  |
| 0.571 | -0.570 | -0.073 | 0.117 | LT (mother) + RF (child) |  |  |
| 0.827 | 0.219 | 0.028 | 0.111 | LT (mother) + LF (child) |  |  |
| 0.973 | -0.033 | -0.004 | 0.112 | LT (mother) + RC (child) |  |  |
| 0.592 | 0.538 | 0.069 | 0.115 | LT (mother) + LC (child) |  |  |
| 0.158 | -1.428 | -0.181 | 0.114 | RF (mother) + RT (child) | -124.691 | RF (mother) |
| 0.127 | -1.547 | -0.196 | 0.117 | RF (mother) + LT (child) |  |  |
| 0.654 | -0.451 | -0.058 | 0.117 | RF (mother) + RF (child) |  |  |
| 0.464 | -0.737 | -0.095 | 0.113 | RF (mother) + LF (child) |  |  |
| 0.235 | -1.200 | -0.153 | 0.118 | RF (mother) + RC (child) |  |  |
| 0.268 | -1.117 | -0.143 | 0.115 | RF (mother) + LC (child) |  |  |
| 0.942 | 0.073 | 0.009 | 0.113 | LF (mother) + RT (child) | -125.298 | LF (mother) |
| 0.981 | -0.023 | -0.003 | 0.120 | LF (mother) + LT (child) |  |  |
| 0.450 | -0.760 | -0.098 | 0.111 | LF (mother) + RF (child) |  |  |
| 0.830 | -0.216 | -0.028 | 0.114 | LF (mother) + LF (child) |  |  |
| 0.889 | -0.141 | -0.018 | 0.114 | LF (mother) + RC (child) |  |  |
| 0.568 | 0.575 | 0.074 | 0.118 | LF (mother) + LC (child) |  |  |
| 0.405 | -0.839 | -0.108 | 0.115 | RC (mother) + RT (child) | -127.781 | RC (mother) |
| 0.570 | -0.572 | -0.074 | 0.120 | RC (mother) + LT (child) |  |  |
| 0.807 | -0.246 | -0.032 | 0.119 | RC (mother) + RF (child) |  |  |
| 0.647 | -0.460 | -0.059 | 0.112 | RC (mother) + LF (child) |  |  |
| 0.549 | -0.603 | -0.078 | 0.118 | RC (mother) + RC (child) |  |  |
| 0.190 | -1.325 | -0.169 | 0.119 | RC (mother) + LC (child) |  |  |
| 0.552 | -0.598 | -0.077 | 0.115 | LC (mother) + RT (child) | -128.356 | LC (mother) |
| 0.966 | 0.043 | 0.006 | 0.119 | LC (mother) + LT (child) |  |  |
| 0.494 | -0.687 | -0.088 | 0.111 | LC (mother) + RF (child) |  |  |
| 0.989 | 0.014 | 0.002 | 0.113 | LC (mother) + LF (child) |  |  |
| 0.183 | -1.347 | -0.171 | 0.119 | LC (mother) + RC (child) |  |  |
| 0.908 | 0.116 | 0.015 | 0.114 | LC (mother) + LC (child) |  |  |

**Correlations between Power and Connectivity – Video chat Paradigm – Child's ROIs**

| **p value (uncorrected)** | **t value** | **r'** | **wPLI Connectivity** |  | **power** | **ROI** |
| --- | --- | --- | --- | --- | --- | --- |
| 0.877 | -0.155 | -0.020 | 0.116 | RT (mother) + RT (child) | -119.816 | RT (child) |
| 0.759 | -0.309 | -0.040 | 0.120 | LT (mother) + RT (child) |  |  |
| 0.963 | 0.046 | 0.006 | 0.114 | RF (mother) + RT (child) |  |  |
| 0.750 | -0.320 | -0.041 | 0.113 | LF (mother) + RT (child) |  |  |
| 0.351 | -0.940 | -0.120 | 0.115 | RC (mother) + RT (child) |  |  |
| 0.857 | 0.182 | 0.023 | 0.115 | LC (mother) + RT (child) |  |  |
| 0.794 | 0.263 | 0.034 | 0.120 | RT (mother) + LT (child) | -119.576 | LT (child) |
| 0.892 | 0.136 | 0.018 | 0.116 | LT (mother) + LT (child) |  |  |
| 0.626 | 0.490 | 0.063 | 0.117 | RF (mother) + LT (child) |  |  |
| 0.794 | 0.263 | 0.034 | 0.120 | LF (mother) + LT (child) |  |  |
| 0.403 | 0.842 | 0.108 | 0.120 | RC (mother) + LT (child) |  |  |
| 0.682 | 0.412 | 0.053 | 0.119 | LC (mother) + LT (child) |  |  |
| 0.378 | 0.888 | 0.114 | 0.116 | RT (mother) + RF (child) | -123.250 | RF (child) |
| 0.757 | 0.311 | 0.040 | 0.117 | LT (mother) + RF (child) |  |  |
| 0.119 | 1.584 | 0.200 | 0.117 | RF (mother) + RF (child) |  |  |
| 0.564 | -0.580 | -0.075 | 0.111 | LF (mother) + RF (child) |  |  |
| 0.371 | 0.901 | 0.116 | 0.119 | RC (mother) + RF (child) |  |  |
| 0.554 | 0.595 | 0.077 | 0.111 | LC (mother) + RF (child) |  |  |
| 0.662 | 0.440 | 0.057 | 0.116 | RT (mother) + LF (child) | -123.447 | LF (child) |
| 0.960 | -0.050 | -0.006 | 0.111 | LT (mother) + LF (child) |  |  |
| 0.779 | -0.282 | -0.036 | 0.113 | RF (mother) + LF (child) |  |  |
| 0.795 | -0.261 | -0.034 | 0.114 | LF (mother) + LF (child) |  |  |
| 0.402 | -0.844 | -0.108 | 0.112 | RC (mother) + LF (child) |  |  |
| 0.936 | -0.081 | -0.010 | 0.113 | LC (mother) + LF (child) |  |  |
| 0.014 | 2.523 | 0.310 | 0.118 | RT (mother) + RC (child) | -125.930 | RC (child) |
| 0.384 | 0.877 | 0.113 | 0.112 | LT (mother) + RC (child) |  |  |
| 0.082 | 1.772 | 0.223 | 0.118 | RF (mother) + RC (child) |  |  |
| 0.074 | 1.819 | 0.229 | 0.114 | LF (mother) + RC (child) |  |  |
| 0.108 | 1.634 | 0.206 | 0.118 | RC (mother) + RC (child) |  |  |
| 0.194 | 1.313 | 0.167 | 0.119 | LC (mother) + RC (child) |  |  |
| 0.247 | 1.170 | 0.149 | 0.114 | RT (mother) + LC (child) | -126.529 | LC (child) |
| 0.819 | 0.229 | 0.030 | 0.115 | LT (mother) + LC (child) |  |  |
| 0.449 | 0.763 | 0.098 | 0.115 | RF (mother) + LC (child) |  |  |
| 0.836 | -0.207 | -0.027 | 0.118 | LF (mother) + LC (child) |  |  |
| 0.767 | -0.298 | -0.038 | 0.119 | RC (mother) + LC (child) |  |  |
| 0.709 | 0.375 | 0.048 | 0.114 | LC (mother) + LC (child) |  |  |

**Table S2. Power and Inter-brain connectivity correlations for each ROI in each Paradigms were not significant, indicating that power did not affect inter-brain Connectivity values.** To validate that the observed differences in wPLI inter-brain connectivity are not related to changes in power, beta power spectral density (PSD) was calculated for each ROI of the mother and child and correlated with the inter-brain connectivity of the relevant ROI. PSD was calculated using MNE's implementation of PSD using multitaper, and PSD scores were calculated for each electrode in each paradigm separately. Next, the power of each ROI was calculated as the average of the 3 relevant electrodes comprising each ROI. The power values of each ROI in each condition (face to face/ video chat) were then correlated with each of the wPLI connectivity values observed for the relevant ROI. Before correcting to multiple compressions (a total of 144 different comparisons are reported here), only 3 correlations yielded a significant link between power and inter-brain connectivity (marked with * in the table), with none of these correlations observed for in an ROI combination that resulted in significant inter-brain synchrony in our study. Following a correction to only 36 comparisons for either mother or child, in either face or skype, none of these correlations reached statistical significance. Therefore, the analysis revealed that power had no effect on the inter-brain connectivity values reported in our study.

**Figure S1 - Histogram of Mean Shuffled wPLI Scores Compared to the Real Data Score**

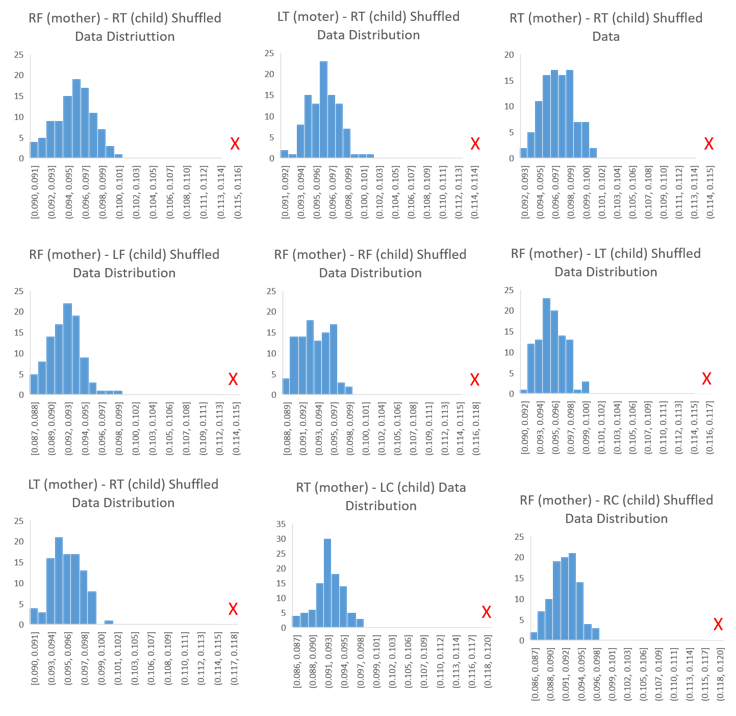

**Figure S1 - Histogram of Mean shuffled wPLI Scores for each of the significant ROIs found in the face to face interactions.** The red 'X' represents the real mean of wPLI scores. For each of the paradigms separately, we randomly shuffled the epochs of one member in each dyad 100 times and compared the original connectivity of each of the real ROI links with the connectivity of values obtained from the shuffled data of the same ROI combinations. For each of the significant links reported in the results section the comparisons the mean of the original data was significantly higher, even after correcting to 36 comparisons for all the data, (nonparametric Wilcoxon signed-rank p (corrected) < 0.001). This analysis was conducted in order to verify that the observed inter-brain neural synchrony is differentiated from spurious synchrony that could be driven by common intrinsic properties of the signal or consistent external perturbation during the experiment.

**Figure S2 – Matching Times across Conditions**

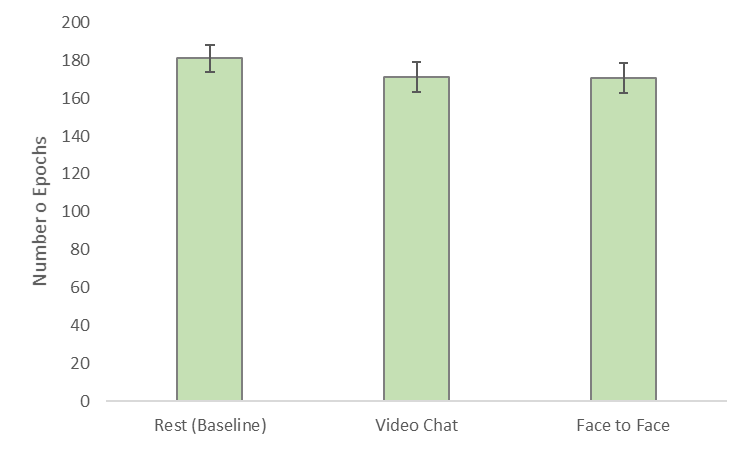

**Figure S2 – Following EEG preprocessing, the number of epochs did not differ across paradigms,** such that only data containing good epochs for both participants was included in the analysis. To compare the inter-subject connectivity scores between two conditions, the interaction duration of each dyad was matched by using the minimal duration among the two for each dyad (Endevelt- Shapira et al, 2021). Following EEG preprocessing and removal of all artifacts, the face-to- face and video chat interactions resulted in the exclusion of surplus epochs compared to the rest condition, for which there were no movement, speech, nor muscular activities and less epochs were excluded. To match the number of epochs in each condition, the following 10 epochs of the face-to-face and video chat interactions were added to the analysis to keep comparability with the number of epochs I the Rest condition and resulted in similar epochs of analysis across all conditions (face to face – M = 170.68, SD = 61.44, video chat = 171.11, SD = 61.68, rest - M = 181.01, SD = 56.11, (F(2,130) = 1.6, p = .21).

**Figure S3 – Significant Inter-brain Connections in each Interaction**

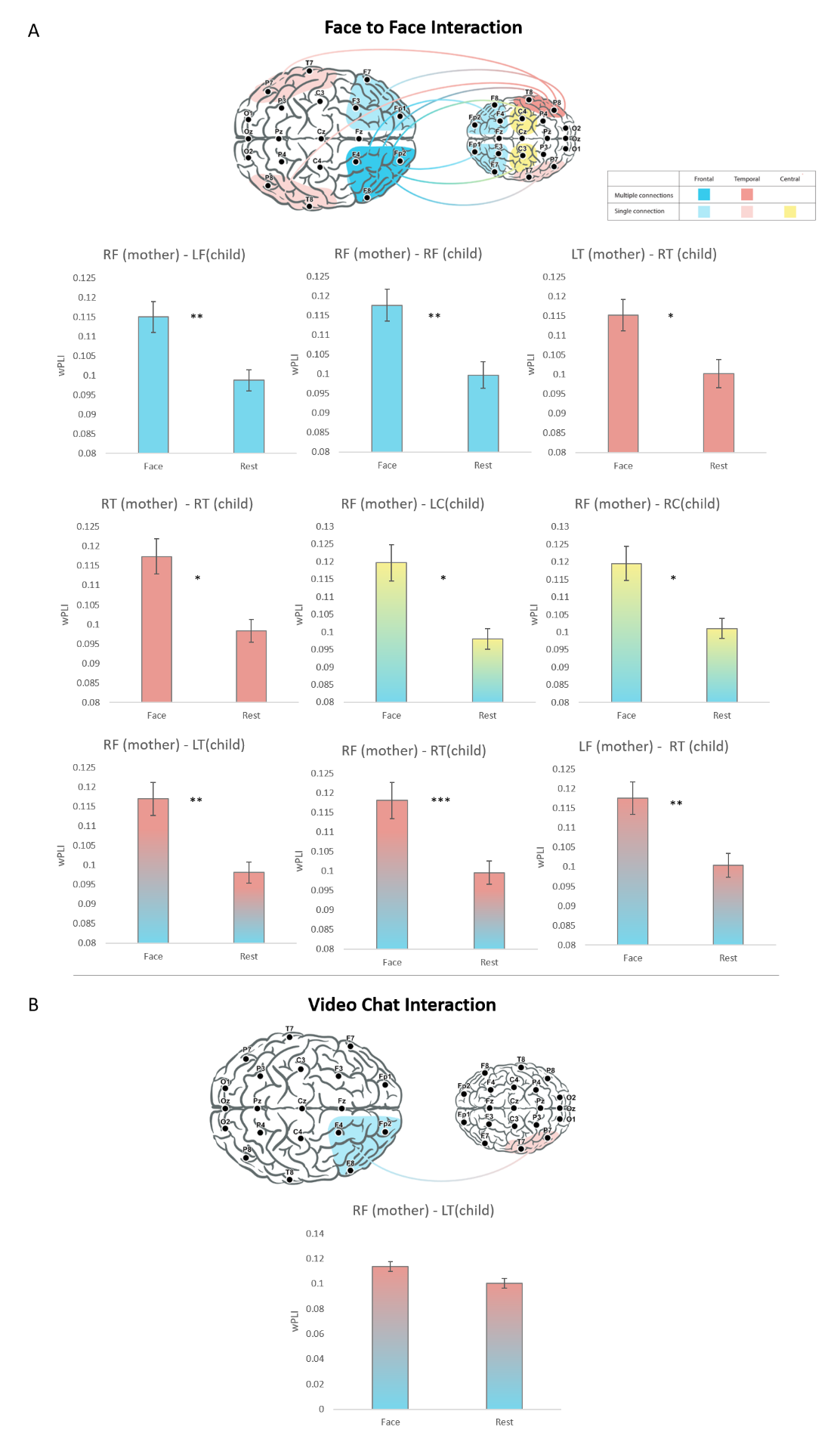

**Fig. S3. (A+ B): Illustration of Mother-child inter-brain neural synchrony in both Face to face and Video chat paradigms**. Inter-brain neural synchrony values were calculated for Beta frequency band (13.5 to 29.5 Hz) using weighted phase lag index (wPLI). 6 regions of interest were pre-defined and examined, each consisting of 3 electrodes. Region of interest were right and left Frontal, Central and Temporal areas (RF, LF, RC, LC, RT, LT, accordingly). Connectivity scores were computed for the 6 regions of interest, resulting in 36 wPLI possible combinations between mother and child synchrony per condition. During face to face interactions 9 inter-brain connections emerged between the mother and child, while in the video chat only one inter-brain connection was found.
